# “Outbreak reconstruction with a slowly evolving multi-host pathogen: a comparative study of three existing methods on *Mycobacterium bovis* outbreaks.”

**DOI:** 10.1101/2023.07.11.548642

**Authors:** Hélène Duault, Benoit Durand, Laetitia Canini

**Author notes:** Corresponding author: (L.C.).

## Abstract

In a multi-host system, understanding host-species contribution to transmission is key to appropriately targeting control and preventive measures. Outbreak reconstruction methods aiming to identify who-infected-whom by combining epidemiological and genetic data could contribute to achieving this goal. However, the majority of these methods remain untested on realistic simulated multi-host data. *Mycobacterium bovis* is a slowly evolving multi-host pathogen and previous studies on outbreaks involving both cattle and wildlife have identified observation biases. Indeed, contrary to cattle, sampling wildlife is difficult. The aim of our study was to evaluate and compare the performances of three existing outbreak reconstruction methods (seqTrack, *outbreaker2* and *TransPhylo*) on *M. bovis* multi-host data simulated with and without biases.

Extending an existing transmission model, we simulated 30 bTB outbreaks involving cattle, badgers and wild boars and defined six sampling schemes mimicking observation biases. We estimated general and specific to multi-host systems epidemiological indicators. We tested four alternative transmission scenarios changing the mutation rate or the composition of the epidemiological system. The reconstruction of who-infected-whom was sensitive to the mutation rate and seqTrack reconstructed prolific super-spreaders. *TransPhylo* and *outbreaker2* poorly estimated the contribution of each host-species and could not reconstruct the presence of a dead-end epidemiological host. However, the host-species of cattle (but not badger) index cases was correctly reconstructed by seqTrack and *outbreaker2*. These two specific indicators improved when considering an observation bias.

We found an overall poor performance for the three methods on simulated biased and unbiased bTB data. This seemed partly attributable to the low evolutionary rate characteristic of *M. bovis* leading to insufficient genetic information, but also to the complexity of the simulated multi-host system. This study highlights the importance of an integrated approach and the need to develop new outbreak reconstruction methods adapted to complex epidemiological systems and tested on realistic multi-host data.

**Author summary:** Some pathogens like the one responsible for bovine tuberculosis can infect multiple species. Identifying which species transmitted and to which other species in such an outbreak presents a unique challenge, especially when difficult to observe wildlife species are concerned. One way to tackle this issue would be to reconstruct who-infected-whom in an outbreak and then identify the role each species played. However, methods that enable this type of reconstruction have not been tested in the context of transmission between unevenly observed species. Moreover, the pathogen responsible for bovine tuberculosis evolves slowly, which further complicates the reconstruction of who-infected-whom. We thus simulated realistic and complex bovine tuberculosis outbreaks on which we tested three widely used methods. We found poor performances for all three tested methods, which highlights the need to develop new methods adapted to outbreaks involving multiple species. Our results also underline the need to combine multiple types of methods and data sources in addition to the reconstruction of who-infected-whom, such as the reconstruction of phylogenetic trees or identifying possible infectious contacts through investigations, when studying an outbreak.

## Introduction

Over 60% of pathogens can infect more than one host-species [1,2]. This possible contribution of multiple host-species to transmission dynamics complicates disease control and surveillance for these multi-host pathogens, especially when one of the host-species to consider is a free-ranging wildlife species. Indeed, quantifying contribution to transmission in order to select appropriate control measures as well as the implementation of said measures can be challenged by the lack of accurate estimations of wildlife population size, the impossibility to restrain the entire wildlife population and the difficulty to prevent interactions between host-species [3]. Multi-host pathogens can have important consequences on human health (*e.g.* zoonotic diseases endemic in wildlife [4]), biodiversity (*e.g.* canine distemper in lions, *Panthera leo*, in the Serengeti national park [5]) and animal trade economy (*e.g.* foot-and-mouth disease and avian influenza [6]).

A prime example of a multi-host pathogen, for which the contribution of wildlife species needs to be considered, is *Mycobacterium bovis*, the most frequent etiological agent of bovine tuberculosis (bTB). Indeed, while *M. bovis* mainly affects cattle, which have been the target of bTB control programs in the European Union since 1964 (EU directive 64/432/EEC), other domestic and wildlife host-species can also be infected [7]. Furthermore, wildlife species have even been implicated around the world as bTB reservoirs, *e.g.* badgers (*Meles meles*) in the United Kingdom [8], wild boars (*Sus scrofa*) in Spain [9] and brush-tailed possums (*Trichosurus vulpecula*) in New Zealand (10). In France, infected wildlife presenting the same genotypes as nearby infected cattle have been reported by the wildlife surveillance program since its implementation in 2012 [11], which suggests bTB transmission between wildlife and cattle and therefore, the presence of bTB multi-host systems.

Studies have aimed to reconstruct phylogenetic trees from *M. bovis* whole genome sequences, present in cattle and wildlife, in order to better understand transmission within these multi-host systems [12–15]. In a phylogenetic tree, internal nodes correspond to hypothetical common ancestors and, using Bayesian methods, the ancestral state (*e.g.* host-species [16,17] or geographical location [18,19]) of these internal nodes can be estimated. These Bayesian methods can therefore reconstruct the host-species of the most recent common ancestor of all sampled sequences [20] as well as transitions between species or groups of individuals over time [15,17], but not transmission events at an individual level. Phylogenetic trees thus differ from transmission trees, in which each node represents an infected host and these infected hosts are linked by directed edges representing transmission events [21]. Such a reconstruction of who-infected-whom in the outbreak makes it possible to estimate transmission parameters specific to each host-species (such as the number of transmission events due to an individual of a particular host-species), and thus sheds more light on the transmission dynamics within the studied multi-host system.

In principle, outbreak (here meaning transmission tree) reconstruction could be based solely on epidemiological data obtained via contact tracing methods (*e.g.* [22]); however data collected are not always reliable nor detailed enough to enable accurate reconstruction [23]. Therefore, some outbreak reconstruction methods have combined both genomic and epidemiological data in transmission tree inference [24–29]. These outbreak reconstruction methods can be divided into two categories according to how genomic data is treated [30], those that consider a link between phylogenetic and transmission trees (generally by annotating branches or internal nodes with infected hosts) [26,28,31,32] and those that solely consider genetic distances [21,25,33]. While some outbreak reconstruction methods were developed to study pathogen transmission within a specific multi-host system (*e.g.* foot-and-mouth disease [34]), most were developed using the example of a single-host system, *e.g.* slowly evolving *M. tuberculosis* [26,31], and more rapidly evolving pathogens like methicillin-resistant *Staphylococcus aureus* [26,33] or SARS-CoV-1 [25] in a human population. However, the development of outbreak reconstruction methods on single-host systems does not preclude them from yielding insightful results in multi-host systems; for instance Willgert *et al.* recently reconstructed the transmission history of SARS-CoV-2 in a human-deer system in Iowa (USA) [35]. In a multi-host system, other than correctly reconstructing transmission events between individuals and estimating outbreak size (general epidemiological indicators), we expect outbreak reconstruction methods to allow accurate estimation of host-species contribution to the outbreak and to identify the host-species of the index case (specific multi-host epidemiological indicators).

While some outbreak reconstruction methods assume that all cases are known and sampled [21,28,36], others account for the presence of unsampled cases by either allowing the annotation of unsampled hosts in the phylogenetic tree [31] or the presence of intermediary unsampled hosts between two sampled hosts [24,25]. When not all cases are sampled in the outbreak, there exists a difference between the actual outbreak and the transmission tree these methods can aim to reconstruct from sampled sequences. Indeed, even if the outbreak reconstruction method accounts for the presence of unsampled hosts [25,31], these hosts can only be inferred if they have descendant sampled hosts [35] and the transmission tree that can be reconstructed is therefore a subtree induced by the sampling process.

The sampling process in a multi-host system that implicates a free-ranging wildlife species can also result in incomplete or even biased data, when observation efforts differ between host-species. For instance, *M. bovis* wildlife surveillance in France was implemented later than cattle surveillance (2012 *vs.* 1954) and only investigates bTB infection in badgers, boars, red deer (*Cervus elaphus*) and roe deer (*Capreolus capreolus*) [11]. However, estimations of bTB infection rates in red foxes (*Vulpes vulpes*) have recently been investigated in France and yielded similar results to those found in badgers and wild boars [37]. These sampling biases between host-species could have an important impact on outbreak reconstruction.

Our aim was to evaluate and compare performances of existing outbreak reconstruction methods on bTB outbreaks in a multi-host system and study whether these performances were affected by sampling biases. Therefore, we simulated bTB transmission within a multi-host system situated in a previously studied area in the South-West of France. In this area, bTB surveillance has reported *M. bovis* circulation in cattle, badgers and wild boars [38]. Multiple sampling schemes were implemented to reflect the late implementation of wildlife surveillance (temporal bias) and the fact that not all host-species are surveilled (species bias). In order to evaluate the quality of reconstructed transmission trees, we calculated general as well as specific multi-host epidemiological indicators.

## Materials and methods

### 1. Reference transmission trees

#### 1.1 Transmission model

We extended an existing model that simulated bTB transmission trees, for the 11 genotypes identified, in a badger-cattle system present in a study area in the South-West of France, from January 2007 to January 2020 [39]. We narrowed our study to one of the two genotypes of *M. bovis*, which were isolated in both wildlife and cattle within our study region. Since infected wild boars have also been detected in this study area [40,41] and our aim was to study a complex multi-host system, we added a wild boar meta-population to the modeled epidemiological system (see details in S1 Appendix). Similarly to the badger population, wild boars could either be susceptible (S) or infected (I) while cattle had an additional latent state (E), when animals could be detected infected but could not transmit the pathogen [39].

Moreover, transmission trees simulated with the original model considered cattle farms and badger social groups as epidemiological units whereas we aimed to reconstruct individual transmission links. Therefore, we extended the model to randomly select infected animals within these groups according to the SEI/SI system dynamics and thus, simulated animal-to-animal transmission. The resulting transmission trees are termed below *reference transmission trees* (terms written in italic are defined in Table 1).

**Table 1.**
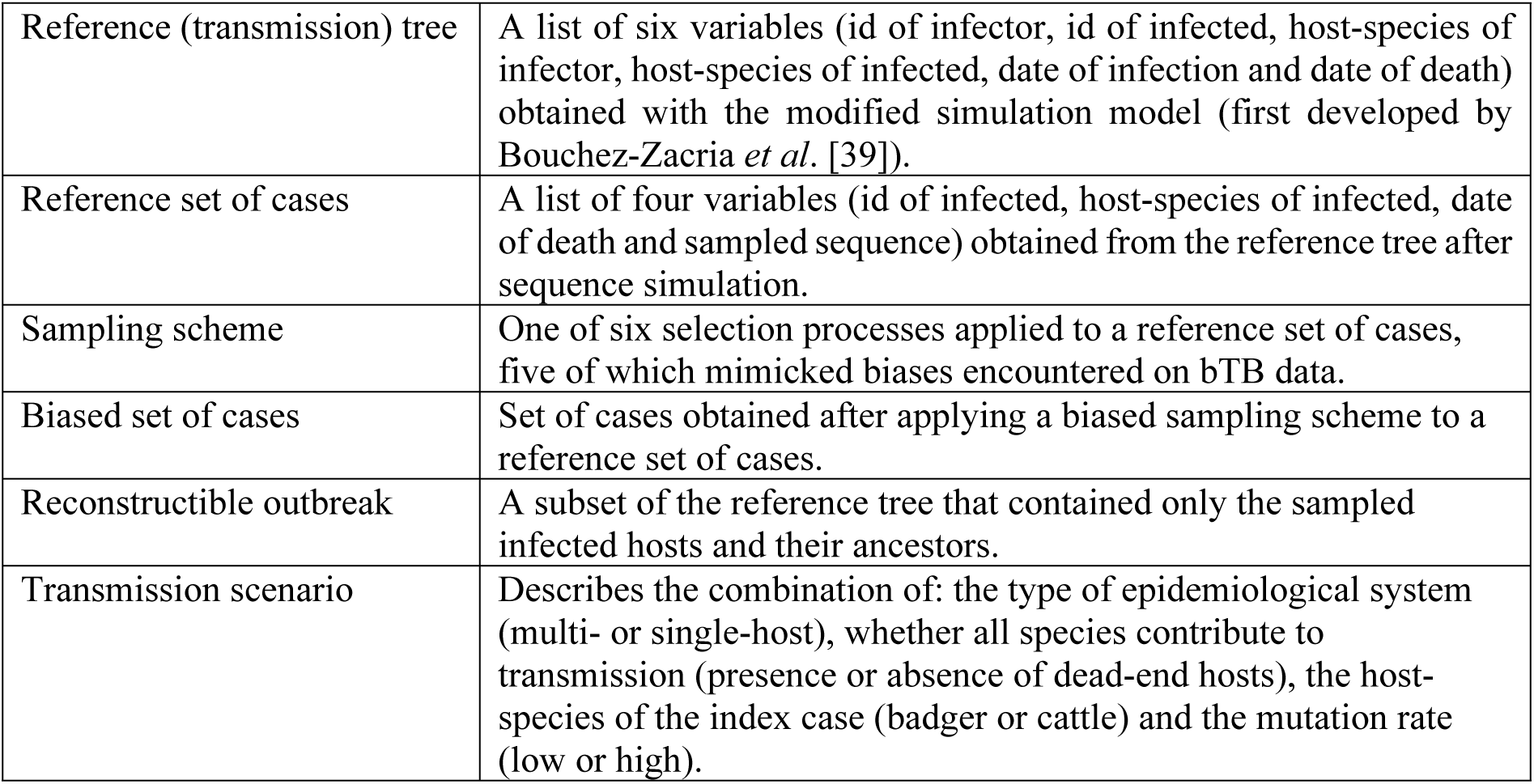
Definition of terms used in the study (in order of appearance in the material and method).

#### 1.2 Reference set of cases

We chose cattle as index cases and bTB spread in the multi-host system was simulated during 13 years. We generated 30 reference transmission trees, in order to investigate various simulated outbreaks while limiting the computational time. These 30 trees had to include less than 500 infected hosts in total, for computational reasons, and at least 15 infected hosts from each host-species, in order to be able to implement sampling schemes. A reference transmission tree corresponded to a list of six variables: identification (id) of infector, id of infected, host-species of infector, host-species of infected, date of infection and date of death.

We simulated genetic sequences along the reference trees according to a Hasegawa-Kishino-Yano (HKY) substitution model (with transition/transversion ratio parameter, κ) [42], since this substitution model was previously used to study *M. bovis* phylogenies [12–14], as well as a fixed mutation rate (µ). We chose µ equal to 0.0024 substitutions per site per year and κ equal to 5.9. Indeed, these values had been previously estimated on 167 *M. bovis* sequences (171 SNPs in length) isolated in cattle and wildlife from this study area [12].

At t = 0, we considered that the index case was infected by a single sequence randomly selected from the 167 sequences isolated in our study area [12]. Our substitution algorithm was based on the Gillespie approach [43] implemented in the *phastSim* package [44] (Fig 1). Taking into account the low genetic diversity observed in *M. bovis* sequences from the same region, we assumed no within-host diversity by considering a single lineage per host but we allowed within-host mutation.

**Fig 1.**
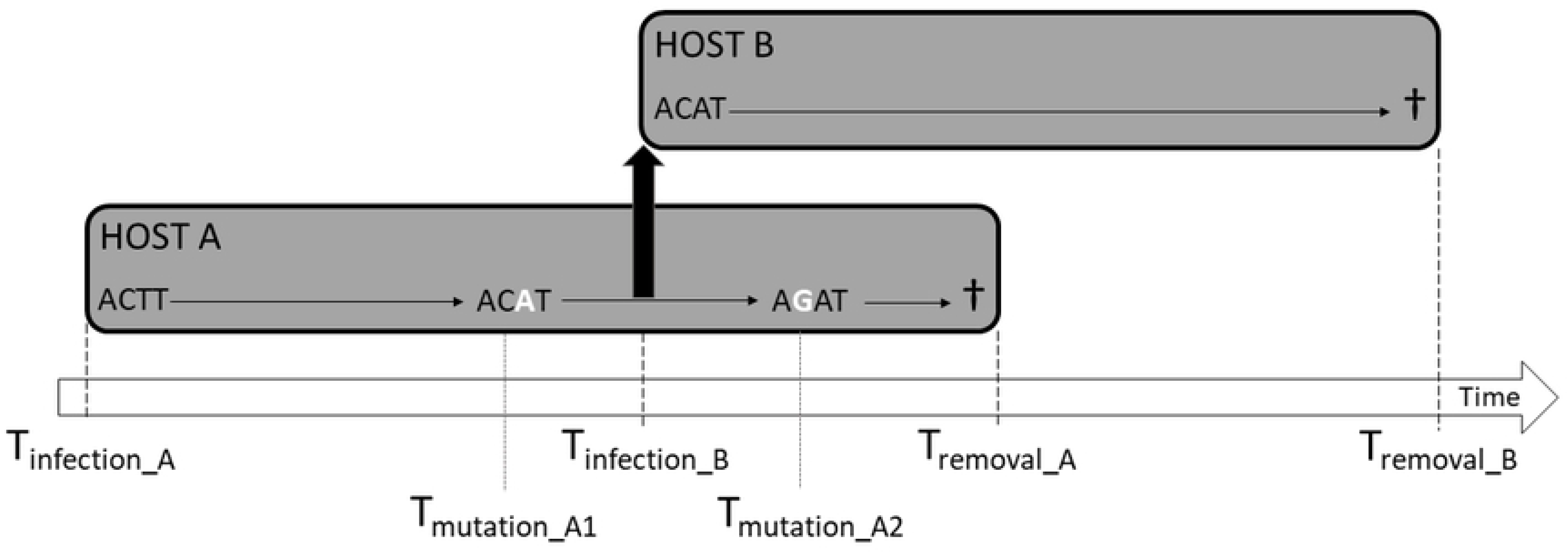
Sequence simulation procedure in two infected hosts A and B. Host A, represented by the grey rectangle on the left (infected at T_infection_A_), transmitted the pathogen to host B at T_infection_B_. This transmission event is represented by the thick black arrow. Hosts were removed (represented by the cross) respectively at T_removal_A_ and T_removal_B_. If the mutation time was inferior to the host removal time (which was the case for T_mutation_A1_ and T_mutation_A2_ in host A), we then selected the nucleotide to mutate (the 3^rd^ nucleotide for the first mutation and the 2^nd^ nucleotide for the second mutation in host A, shown in white) and changed it according to a substitution model. If the mutation time was superior to the removal time of the host (see host B), the sequence did not change until host removal and this sequence was then the one sampled from the host.

We simulated sequences until February 2020. Then, the last simulated sequence was recorded for each host, which corresponded to either the sequence present at the time of removal or in February 2020, for infected hosts not yet removed at the end of the simulation. For each reference transmission tree, we thus obtained a *reference set of cases* (Table 1), meaning a list of four variables: id of infected, host-species of infected, date of death (or February 2020 if host still alive) and sampled sequence.

### 2. Sampling schemes and reconstructed transmission trees

#### 2.1 Sampling schemes

We first considered the hypothetical situation where all infected hosts are observed (reference *sampling scheme*, Table 1), which corresponds to the reference set of cases. Then, we simulated five sampling schemes that mimicked observation biases in bTB epidemiological data, while also sampling all infected hosts unaffected by the scheme (even those not yet removed at the end of the simulation). In scheme T (for “temporal bias”), the late implementation of wildlife surveillance in the study region was simulated and we only considered wildlife cases after 2012. Moreover, the fact that not all host-species are surveilled was simulated in schemes S (for “species bias”): either wild boar cases were not considered, scheme S_W_, or badger cases, in scheme S_B_. Finally, in scheme T+S_W_ (or T+S_B_), we disregarded cases before 2012 for the remaining wildlife species (respectively badgers and wild boars).

We thus simulated for each reference transmission tree one *biased set of cases* (Table 1) for each sampling scheme (T, S_B_, S_W_, T+S_B_, T+S_W_), that contained the same variables as the reference set of cases. With 30 reference trees for each sampling scheme, we thus obtained a total of 30*6 = 180 sets of cases. For each of these sets of cases, we extracted from the reference transmission tree, the *reconstructible outbreak* (Table 1), which is the subtree containing only the cases that were sampled and their ancestors.

#### 2.2 Transmission tree reconstruction

From our review on outbreak reconstruction methods [30], we identified three methods (seqTrack, *outbreaker2* and *TransPhylo*) that were available in an R package and that needed only sampling and/or removal times. In seqTrack and *outbreaker2*, transmission is estimated based on pairwise genetic distances, while in *TransPhylo,* a link is established between phylogenetic and transmission trees [30].

○ seqTrack

Using Edmonds’ algorithm, seqTrack computes the transmission tree in which the total genetic distance between nodes is minimal, assuming that infectors are sampled before the host they infected [21]. In order to use this method, we estimated pairwise genetic distances by using the dist.dna function (*ape* R package v.5.4-1 [45]) with the F84 substitution model since it closely resembles the HKY model [42]. seqTrack [21] is a function available in the *adegenet* R package [46,47]. The format of the tree reconstructed by seqTrack was a table with five columns corresponding to the following variables: id (indices of infected hosts), ances (indices of infectors), weight (number of mutations separating infected hosts from their infectors), date (sampling date of the infected host), ances.date (sampling date of their infector).

○ *outbreaker2*

*outbreaker2* is a Bayesian method that considers four likelihoods: genetic, temporal, reporting and contact [25]. In this method, probability of transmission is inferred from known generation time (time between the infection of a case and the time of transmission from that case to secondary cases) and sampling interval (time from infection to sampling) distributions. Here, we assumed that generation time and sampling interval nonparametric distributions could be obtained without bias by estimating them from the reference trees, which contained every infected host, timed transmission event between hosts and host sampling time. We selected a chain length of 100,000 iterations, a sampling frequency of 1 in 50 and a burn-in period of 10% (for details on priors used and other arguments see S1 Appendix). We graphically checked for convergence and independence of sampling (Effective Sample Size (ESS) above 200 for each parameter), after estimation using the *coda* R package v.0.19-4 [48]. When the ESS were lower than 200, we ran an additional 100,000 iterations and then checked the ESS again. This step was repeated until every ESS was above 200.

Then, we built the consensus tree, as suggested by the authors, computing the most frequent infector for each infected host in the posterior trees as well as the support (posterior probability) for each transmission event. By construction, cycles can be present in this consensus tree (which then becomes a directed graph), meaning that infected hosts can be both the ancestors and the descendants of other infected hosts. Moreover, since this method considers a reporting likelihood, the probability of sampling an infected host is estimated and unsampled hosts are indirectly represented in the consensus tree, as a number of generations separating two sampled hosts.

The format of the consensus tree reconstructed by *outbreaker2* was a table with five columns corresponding to the following variables: from (indices of infectors), to (indices of infected hosts), support (transmission probability), time (estimated time of transmission), date (sampling date of the infected host) and generations (number of intermediary hosts + 1).

○ *TransPhylo*

*TransPhylo*, another Bayesian method, affects infected hosts along branches in a previously reconstructed phylogenetic tree [31] (for details on phylogenetic reconstruction see S1 Appendix). We assumed that the generation time and sampling interval followed a Gamma distribution and that the mean and standard deviation could be obtained without bias by estimating them from the reference trees using the *epitrix* R package v.0.2.2 [49]. We selected a number of iterations of 500,000, a sampling frequency of 1 in 50 and a burn-in period of 20% (for details on priors used and other arguments see S1 Appendix). We used the same method as with *outbreaker2* to check for convergence and independence of sampling, however we considered a lower threshold for the ESS, 100 for each parameter as suggested by the authors [50]. When the ESS were lower than 100, we ran an additional 500,000 iterations and then checked the ESS again. This step was repeated until convergence and independence of sampling parameters were satisfied or the number of iterations reached 2,500,000, we then discarded the reference trees for which convergence was not obtained in every sampling scheme.

Then, as described by *Didelot et al.* [50], we computed the medoid transmission tree (the transmission tree that is the least different from all other posterior trees according to a distance metric defined by Kendall *et al.* [51]). This method accounts for the presence of unsampled hosts when affecting hosts to branches in the phylogenetic tree, and unsampled hosts are explicitly represented as nodes in the medoid transmission tree. This means that in the medoid tree, contrary to the consensus tree in *outbreaker2*, unknown infected hosts can be responsible for more than one transmission event. As in *outbreaker2*, *TransPhylo* estimates a sampling probability.

The format of the medoid tree reconstructed by *TransPhylo* was a table with four columns corresponding to the following variables: tinfection (estimated time of infection), tremoved (estimated time of removal of the infected host), infector_id (id of infector), infected_id (id of infected).

From the sampled posterior trees, we also computed the n-by-n matrix of transmission probability using the computeMatWIW function implemented in *TransPhylo*, where n is the number of sampled infected hosts. Then, we identified for each infected host, its most likely infector corresponding to the infector with the highest probability in the matrix of transmission probabilities. If this probability was zero, we considered the most likely infector of the infected host to be unknown. Note that this method of summarizing posterior trees can lead to the presence of cycles, as in *outbreaker2*, and since time of infection is not estimated, no index case can be inferred.

### 3. Genetic information and epidemiological indicators

#### 3.1 Genetic information

To understand the impact of the sequence simulation model on outbreak reconstruction and facilitate comparison with other works, we first quantified the genetic diversity present in each simulated set of cases. We estimated the proportion of unique sequences in every set of cases obtained with the reference sampling scheme as well as the mean transmission divergence. Transmission divergence was defined in Campbell *et al.*’s work [52] as the number of SNPs separating known transmission pairs, we used reference transmission trees to identify transmission pairs and calculated the mean number of SNPs separating these transmission pairs for every reference tree.

#### 3.2 Epidemiological indicators

*TransPhylo* had two different outputs (the medoid tree and transmission probability matrix). We used the transmission probability matrix when evaluating the method’s accuracy and the medoid tree for all other indicators.

○ Accuracy

In order to evaluate the performance of all three reconstruction methods, we first determined the correct transmission events that could be reconstructed between individuals from each simulated set of cases. For the reference set of cases, the correct transmission events were those present in the reference trees. However, for each biased set of cases, we considered that the correct transmission events were those that connected observed cases to each other, bypassing intermediary unobserved cases. For instance, the chain of transmission Sampled subject #1 → Unobserved subject #2 → Sampled subject #3 would become Sampled subject #1 → Sampled subject #3. For all three methods, we estimated whether reconstructed infector-infected pairs (meaning every “id”-“ances” for seqTrack, “from”-“to” for *outbreaker2* and “infector_id”-“infected_id” for the transmission matrix estimated from *TransPhylo*) were one of the correct transmission events or not.

○ Presence of super-spreaders

For all three methods, we considered super-spreaders to be present in a reconstructed tree when less than 10% of infected hosts were responsible for over 80% of transmission events. Moreover, when super-spreaders were present in a reconstructed tree, we identified the maximum number of transmission events a single super-spreader could be responsible for as well as the host-species of said super-spreader.

○ Host-species of the index case

We evaluated the ability of all three methods to reconstruct the correct host-species of the index case (*i.e.* cattle). Contrary to the *TransPhylo* medoid trees, in which identifying the index case is straightforward (“infected_id” with the earliest “tinfection”), the presence of cycles in *outbreaker2* and the multiples index cases possible in seqTrack complicated the identification of the index case. For seqTrack, we considered the most frequent host-species from the reconstructed index cases (“id” for whom the “ances” is unknown). For *outbreaker2*, we considered the host-species of the index case to be the most frequent host-species among cases infected at the earliest date (“to” with the earliest “time”).

○ Outbreak size

We evaluated the ability of *outbreaker2* and *TransPhylo* to estimate the size of the outbreak (seqTrack does not estimate outbreak size and was thus excluded for this indicator). The simulated outbreak size was the number of infected hosts present in each reference tree. We calculated the corresponding estimate by dividing the number of sampled hosts in each reconstructed tree with the median of the sampling proportion provided by *outbreaker2* and *TransPhylo.* In addition, we tested if the results for this indicator differed depending on whether we were considering the reconstructible outbreak or the reference tree. Therefore, we also calculated the number of infected hosts present in the reconstructible outbreak, and compared it with the number of hosts (sampled and unsampled) present in the trees reconstructed by *outbreaker2* and *TransPhylo*.

○ Host-species contribution

Considering the importance of identifying the host-species that contributed the most to transmission in a multi-host system, we evaluated the ability of *outbreaker2* and *TransPhylo* to reconstruct the number of transmission events due to each host-species. Similarly to the outbreak size, seqTrack was also excluded. The number of transmission events due to each host-species was first calculated in the reference trees. As for the outbreak size, we calculated the corresponding estimate by dividing the number of transmission events between sampled hosts in each reconstructed tree with the median of the sampling proportion provided by *outbreaker2* and *TransPhylo*. We then calculated the number of transmission events due to each host-species in the reconstructible outbreak. This number was compared to the number of all transmission events (to sampled and unsampled infected hosts) due to each host-species present in the reconstructed trees.

○ Statistical analysis

For the outbreak size and host-species contribution estimates, we obtained a credible interval using the bounds of the 95%HPD (High Posterior Density) interval. For each reconstructed tree, we evaluated whether the credible interval contained the simulated outbreak size or number of transmission events due to each host-species. For all epidemiological indicators except the presence of super-spreaders, we tested the effect on the indicator value of the outbreak reconstruction method as well as its interaction with the effect of sampling scheme. In order to account for the non-independence of reconstructed trees (six sets of cases are constructed from the same reference tree), we fit mixed-effects models, using the id of the reference tree as a random effect. For accuracy and index case, we selected a binomial distribution and the probability of either reconstructing a correct transmission event or the correct host-species for the index case was set as the outcome. Due to the overdispersion present in the estimates of outbreak size and number of transmission events, we considered for both indicators a negative binomial distribution. Since, for outbreak size and host-species contribution, we aimed to compare estimates with the values in either the reference tree or the reconstructible outbreak, these values were set as an offset and the intercept was set to zero. The estimated incidence rates ratios (IRRs) could therefore be interpreted as multiplicative factors of the outbreak size (or host contribution) in the reference tree (or reconstructible outbreak).

### 4. Alternative transmission scenarios

We tested the influence of the low evolutionary rate, which is characteristic of *M. bovis*, on our results. We simulated new sequences along the 30 reference trees having increased the mutation rate by a factor of 10 (µ_h_ = 0.024 substitutions per site per year) and implemented the three outbreak reconstruction methods on the reference set of cases only.

To test whether the reconstruction of outbreak size and accuracy were influenced by the complexity of the epidemiological system, we then simulated 30 new reference trees of a single-host system, by setting transmission parameters to, between and from wildlife to 0, in order to obtain cattle-only transmission trees. We simulated sequences along these 30 new trees with µ (0.0024 substitutions per site per year), then implemented the three methods on these sequences.

We then analyzed whether asymmetrical roles within the multi-host system influenced the reconstruction of the host-species contributions. With the same protocol (30 reference trees and a low evolutionary rate), we tested a transmission scenario where one of the host-species could be infected but could not play any role in transmission (dead-end epidemiological host). We obtained a multi-host system where wild boars played no part in onward bTB transmission by setting transmission parameters between and from wild boars to 0.

Finally, in order to evaluate the reconstruction of the host-species of the index case, we simulated 30 new reference trees with badgers as index cases, in the multi-host system where every host-species contributed to transmission.

## Results

### 1. Transmission tree reconstruction

While convergence was not a limiting factor for *outbreaker2*, it could not be obtained for every set of cases in BEAST2 nor every consensus phylogenetic tree with *TransPhylo*. We were thus restrained to 21 out of 30 reference trees (126 reconstructed trees in total). The reference trees from which we could not reconstruct trees in *TransPhylo* showed a higher median number of infected hosts compared to those whose set of sequences and trees converged (S1 Table).

Computational time varied greatly between sets of cases (or consensus phylogenetic trees) and reconstruction methods: less than 10 min for all 126 trees reconstructed by seqTrack, from less than 20 min (when only 100,000 iterations were needed) to two hours per tree reconstructed by *outbreaker2*, and from less than an hour to over 12 hours (for 2,500,000 iterations) per tree reconstructed by *TransPhylo*. Moreover, phylogenetic reconstruction with BEAST2 was needed to implement *TransPhylo* and computational time also varied between sets of cases: from five hours to two days. In total, the computational time for these 378 (126 trees*3 methods) reconstructed trees was around three months.

The median proportion of unique sequences in the reference set of cases for which convergence was obtained was 6.1%. The median of the mean transmission divergence was 0.19 (S1 Table) and the majority of transmission pairs shared the same sequence (S1 Fig).

All trees reconstructed by *outbreaker2* as well as all transmission probability matrices estimated by *TransPhylo,* for which we kept the most probable infectors, presented cycles.

### 2. Epidemiological indicators

#### 2.1 Accuracy

When all sequences were sampled, the median proportion of correctly reconstructed transmission events (Fig 2) was 3.4% (range: 1.3-12.1) for trees reconstructed by seqTrack, 8.0% (2.2-11.3) for *outbreaker2* and 8.9% (6.0-16.8) for *TransPhylo* (S2 Table).

**Fig 2.**
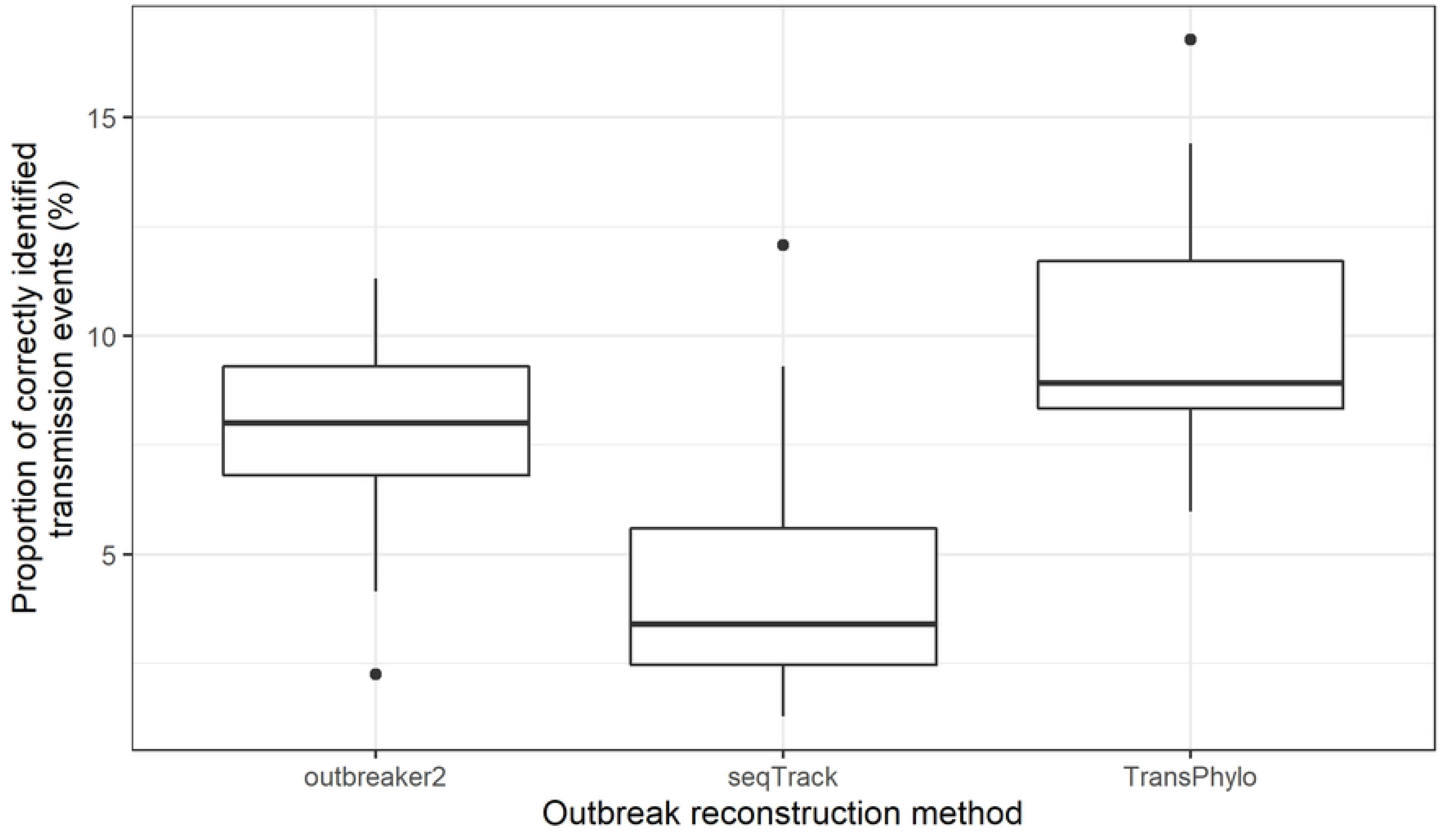
**Proportion of transmission events reconstructed from all sequences present in reference trees according to method.**

Compared to *outbreaker2*, the probability of reconstructing a correct transmission event was significantly lower for seqTrack (OR=0.51, p-value<0.001) but significantly higher for *TransPhylo* (OR=1.30, p-value<0.001) (Table 2). In trees reconstructed by seqTrack, sampling schemes where wild boars were not sampled increased significantly the probability of reconstructing a correct transmission event (OR=1.37 and 1.30, p-value=0.001 and 0.008 for S_W_ and T+S_W_ respectively). Results did not show a significant effect of the sampling scheme on accuracy for the other two methods (Table 2).

**Table 2.** Presence of reconstructed transmission events in reference trees tested with a Binomial GLMM using reconstruction method, interaction between method and sampling scheme as fixed effects.

| Fixed effects | *outbreaker2* |  | seqTrack |  | *TransPhylo* |  |
| --- | --- | --- | --- | --- | --- | --- |
|  | OR | p-value | OR | p-value | OR | p-value |
| Method | - | - | **0.51** | **<0.001** | **1.30** | **<0.001** |
| Method :T | 0.99 | 0.89 | 0.94 | 0.54 | 0.91 | 0.17 |
| Method :$S_B$ | 0.91 | 0.21 | 0.95 | 0.60 | 1.08 | 0.30 |
| Method :$T+S_B$ | 0.92 | 0.32 | 0.94 | 0.57 | 1.11 | 0.14 |
| Method :$S_W$ | 1.08 | 0.33 | **1.37** | **0.001** | 1.03 | 0.64 |
| Method :$T+S_W$ | 1.07 | 0.41 | **1.30** | **0.008** | 1.01 | 0.88 |

OR stands for odds ratio. Results in bold mean that the p-value was <0.05. The *outbreaker2* method, the reference sampling scheme was set as reference, hence the “-“ present on the method line and the absence of the reference sampling scheme. T stands for “temporal bias”, $S_B$ for “badger bias” and $S_W$ for “wild boar bias”. $T+S_B$ ( $T+S_W$ ) combined the temporal and the badger (wild boar) bias.

#### 2.2 Super-spreaders

While in the reference trees the maximum number of transmission events a single infected host could be responsible for ranged from 9 to 27 (median: 14) and no super-spreaders were identified, all trees reconstructed by seqTrack presented super-spreaders. The median of the maximum number of transmission events a single super-spreader could be responsible for ranged from 90 to 108, while the median number of transmission events in the reconstructed trees ranged from 200 to 244 (S3 Table). The most frequent host-species responsible for this maximum number of transmission events was cattle (from 57% in the reference sampling scheme to 86% in the combined temporal and wild boars bias). None of the trees reconstructed by the two other methods presented super-spreaders.

#### 2.3 Host-species of the index case

When all sequences were sampled, the proportion of correctly reconstructed host-species of the index case (*i.e.* cattle) was 76% for trees reconstructed by seqTrack, 81% for *outbreaker2* and 57% for *TransPhylo* (S4 Table). Except when considering the temporal bias alone with the *TransPhylo* method, a temporal and a badger bias (combined or not) led to an increase in the proportion of correctly reconstructed index cases.

#### 2.4 Outbreak size

In the reference trees, the median number of infected hosts was 245 (S5 Table). Overall, the simulated outbreak size was close to the credible interval estimated by *outbreaker2* (Fig 3). Indeed, this credible interval contained the simulated outbreak size for all 21 trees reconstructed with the reference and temporal sampling scheme. However, a species bias (combined or not with a temporal bias) decreased the number of trees that correctly estimated the outbreak size and led to a majority of trees that underestimated the outbreak size (20/21 with S_B_ and T+S_B_, 16/21 for S_W_ and 18/21 for T+S_W_). According to the statistical model, the outbreak size estimated by *outbreaker2* was not significantly different to the reference tree size (IRR= 1.14, p-value=0.43) and sampling schemes had no significant effect on outbreak size (Table 3).

**Fig 3.**
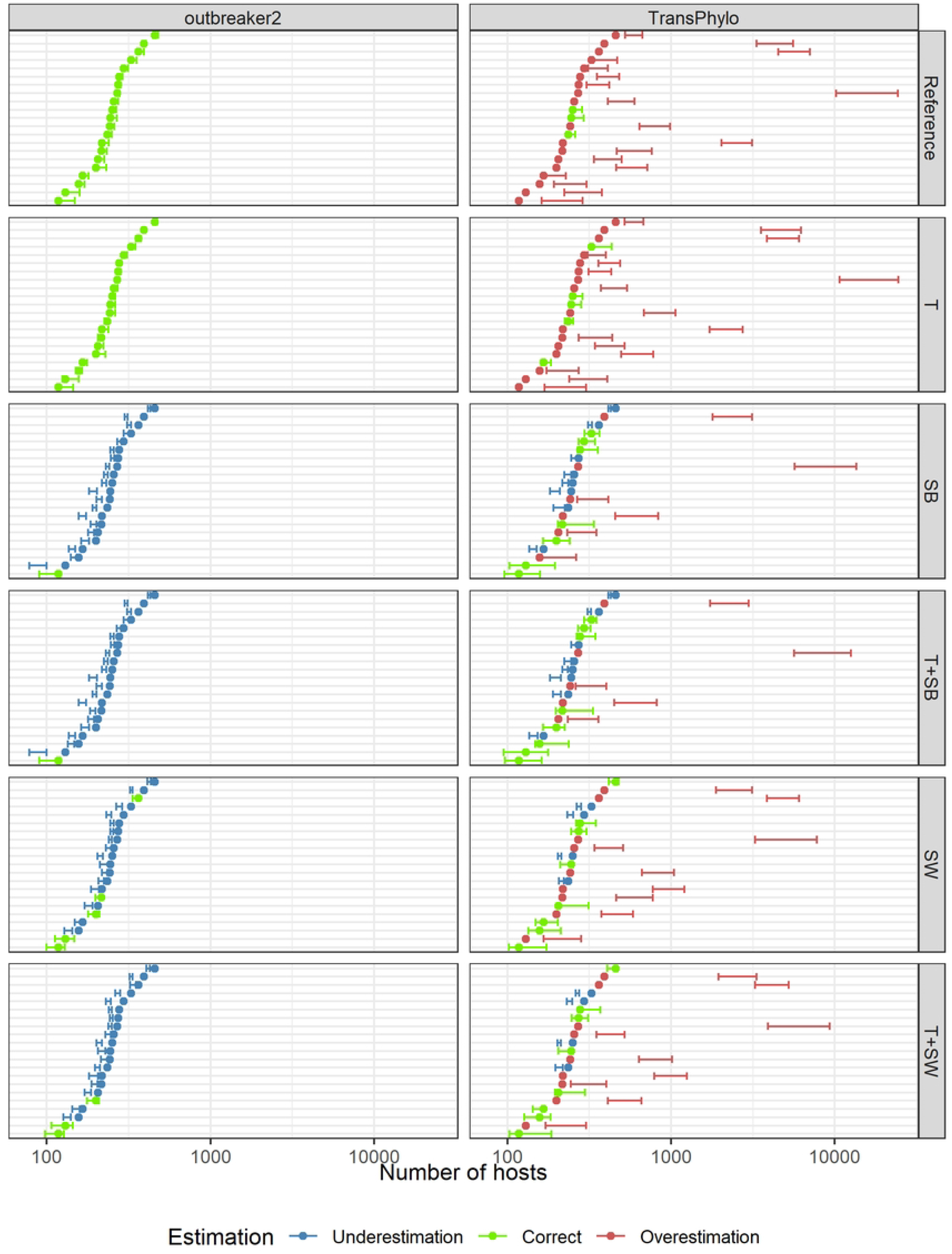
Outbreak size credible interval estimated by *outbreaker2* and *TransPhylo* compared to simulated outbreak size. The point corresponds to the simulated outbreak size. T stands for “temporal bias”, S_B_ for “badger bias” and S_W_ for “wild boar bias”. T+S_B_ (T+S_W_) combined the temporal and the badger (wild boar) bias.

**Table 3.**
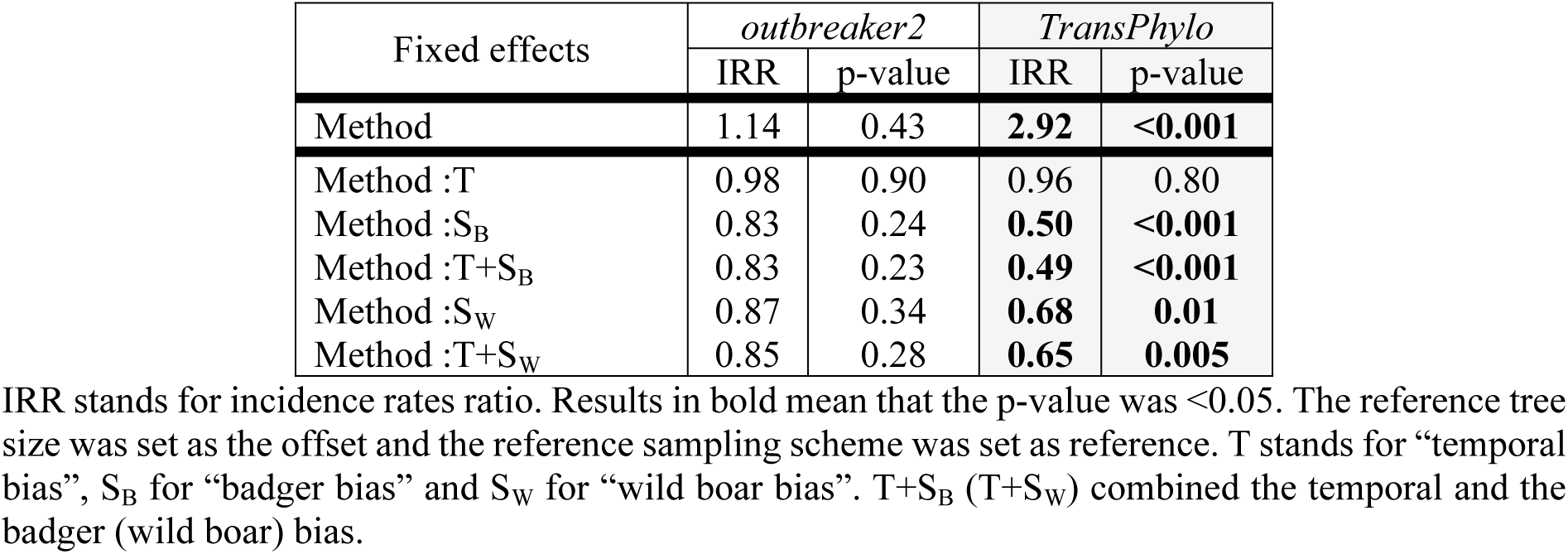
Estimated outbreak size tested with a Negative Binomial GLMM using reconstruction method, interaction between method and sampling scheme as fixed effects.

| Fixed effects | *outbreaker2* |  | *TransPhylo* |  |
| --- | --- | --- | --- | --- |
|  | IRR | p-value | IRR | p-value |
| Method | 1.14 | 0.43 | **2.92** | **<0.001** |
| Method :T | 0.98 | 0.90 | 0.96 | 0.80 |
| Method :S<sub>B</sub> | 0.83 | 0.24 | **0.50** | **<0.001** |
| Method :T+S<sub>B</sub> | 0.83 | 0.23 | **0.49** | **<0.001** |
| Method :S<sub>W</sub> | 0.87 | 0.34 | **0.68** | **0.01** |
| Method :T+S<sub>W</sub> | 0.85 | 0.28 | **0.65** | **0.005** |

IRR stands for incidence rates ratio. Results in bold mean that the p-value was <0.05. The reference tree size was set as the offset and the reference sampling scheme was set as reference. T stands for “temporal bias”, S<sub>B</sub> for “badger bias” and S<sub>W</sub> for “wild boar bias”. T+S<sub>B</sub> (T+S<sub>W</sub>) combined the temporal and the badger (wild boar) bias.

*TransPhylo* could greatly overestimate the outbreak size and the difference between the lower bound of the interval and the simulated outbreak size could exceed 10,000 infected hosts (Fig 3). The reference and temporal sampling scheme led to an overestimation of the outbreak size in the majority of reconstructed trees (19/21 and 16/21): the credible intervals contained the simulated outbreak size in 3 and 5 out of 21 trees, respectively. The number of correct estimations remained low for the other types of biases. The statistical model confirmed these results, since the outbreak size estimated by *TransPhylo* was significantly higher than the simulated outbreak size (IRR= 2.92, p-value <0.001). Moreover, all biased schemes except for the temporal bias significantly lowered the estimated outbreak size (IRR ranging from 0.49 to 0.68 and p-value <0.01).

#### 2.5 Host-species contribution to transmission

The median number of transmission events due to each host-species in the reference trees was 175 for cattle, 24 for badgers and 40 for wild boars (S6 Table).

In the reference sampling scheme, the credible interval contained the simulated number of transmission events due to each host-species in few of the trees reconstructed by *outbreaker2* (2/21 trees for cattle, none for badger and wild boars) and *TransPhylo* (5/21 trees for cattle, 4/21 for badgers and 3/21 for wild boars) (Fig 4). Otherwise, the number of transmission events in the majority of the remaining trees was either underestimated (cattle: 14/21 trees for *outbreaker2* and 13/21 trees for *TransPhylo*), overestimated (badgers: 19/21 trees for *outbreaker2* and 13/21 trees for *TransPhylo*) or no particular trend was observed (wild boars). Similar results were obtained with the other five sampling schemes (S2-S4 Figs).

**Fig 4.**
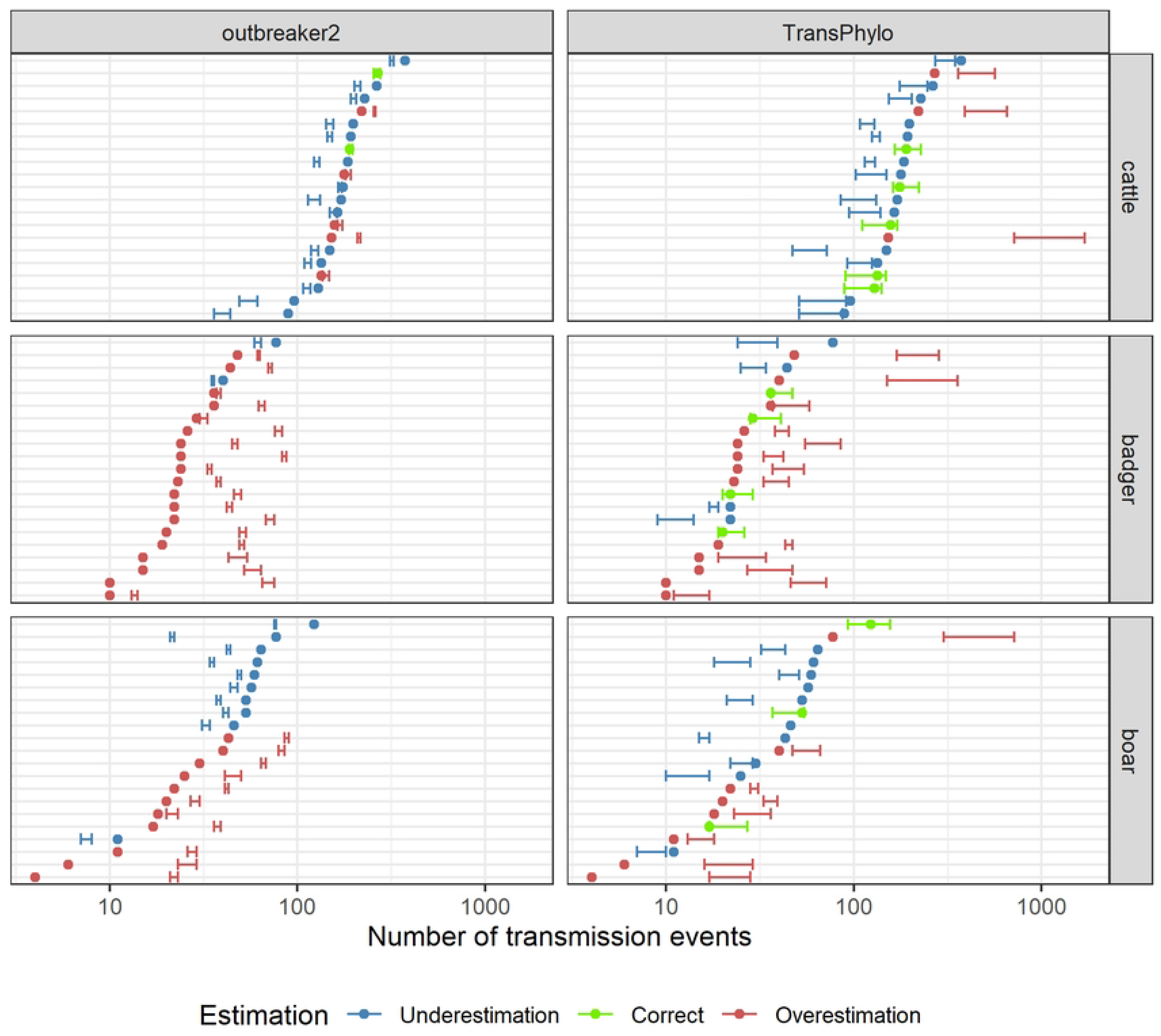
Credible interval of host-species contribution compared to simulated outbreaks. The credible interval was either estimated by *outbreaker2* or by *TransPhylo*. The point corresponds to the number of transmission events due to each host-species in the simulated outbreak. Only the reference sampling scheme is considered here.

According to the statistical model, the underestimation of the number of reconstructed transmission events due to cattle (Fig 4) was not significant for either method (Table 4). The statistical model confirmed the results obtained for badgers, since the number of transmission events due to badgers estimated by both methods was significantly higher than the simulated number (IRR=2.06 for *outbreaker2* and 1.70 for *TransPhylo*, p-value<0.001) (Table 4). Results did not show a significant effect of the sampling scheme on badger contribution for *outbreaker2*. However, the sampling scheme with the least number of sampled hosts (temporal and wild boar biases combined) significantly decreased the number of transmission events due to badgers compared to the reference sampling scheme in trees reconstructed by *TransPhylo*. Finally, the number of transmission events due to wild boars estimated by both methods was not significantly different to the simulated number in the reference tree (Table 4).

**Table 4.**
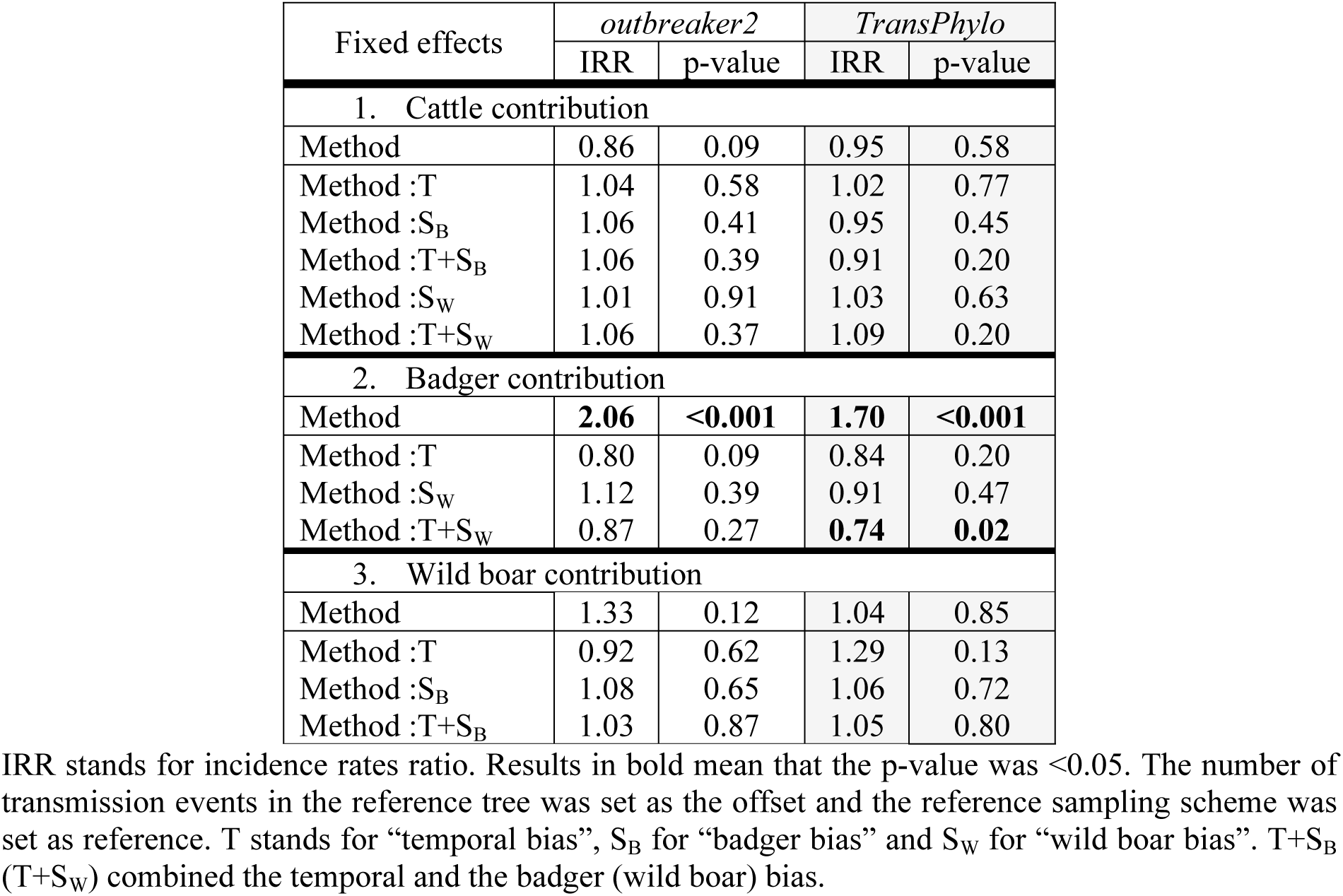
Number of transmission events due to each host-species tested with a Negative Binomial GLMM per host-species using method and interaction between method and sampling scheme as fixed effects.

### 3. Alternative transmission scenarios

#### 3.1 Higher mutation rate

As expected, sequences simulated with a higher mutation rate presented a higher proportion of unique sequences (median: 33.4%) and a higher mean transmission divergence (median: 0.69) (S7 Table).

A higher mutation rate increased markedly the median accuracy for all three methods: 25.7% (+17.7, range: 15.9-33.3) for *outbreaker2*, 15.3% (+11.9, range: 8.2-33.3) for seqTrack and 21.2% (+12.3, range: 13.2-29.3) for *TransPhylo* (S8-S9 Tables). While the majority of trees reconstructed by seqTrack again contained super-spreaders (20/21), the median of the maximum number of transmission events due to a single super-spreader was lower when considering a higher mutation rate (35 *vs.* 108, S10 Table).

The credible interval contained the simulated outbreak size for 16/21 trees reconstructed by *outbreaker2* and in only 4/21 trees reconstructed by *TransPhylo*, otherwise the outbreak size was overestimated (Fig 5 and S11 Table). For both methods, the credible interval contained the number of transmission events due to each host-species in only 4/21 trees for cattle, 3/21 (*outbreaker2*) and 5/21 (*TransPhylo*) for badgers and 1/21 for wild boars (Fig 6). Otherwise, cattle contribution was underestimated by *TransPhylo* (16/21 trees in Fig 6, and S12 Table) and wildlife contribution was overestimated by *outbreaker2* (13/21 trees for badgers and wild boars in Fig 6, and S12 Table).

**Fig 5.**
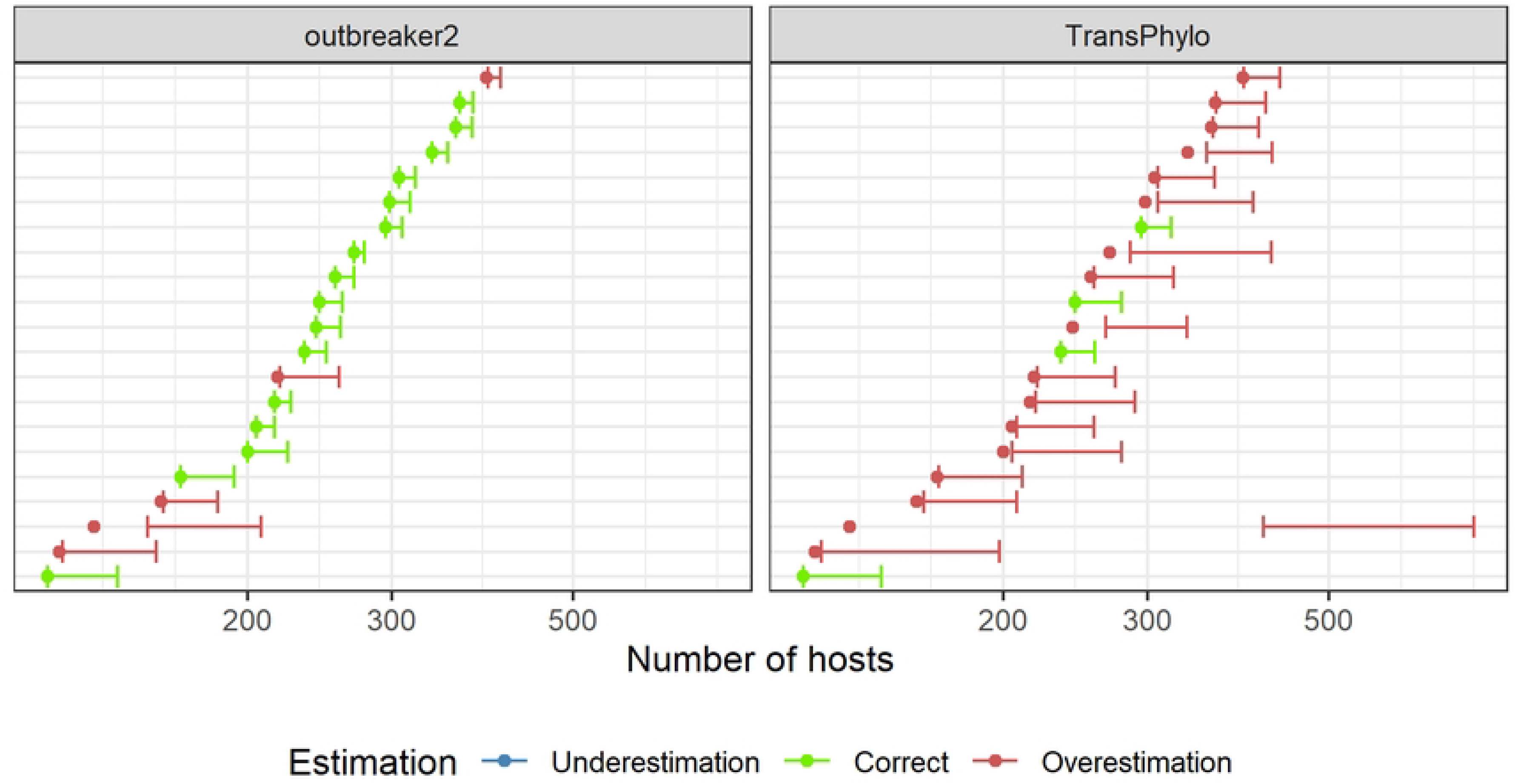
Outbreak size credible interval compared to simulated outbreak size in the high mutation rate scenario. The credible interval was either estimated by *outbreaker2* or by *TransPhylo*. The point corresponds to the simulated outbreak size.

**Fig 6.**
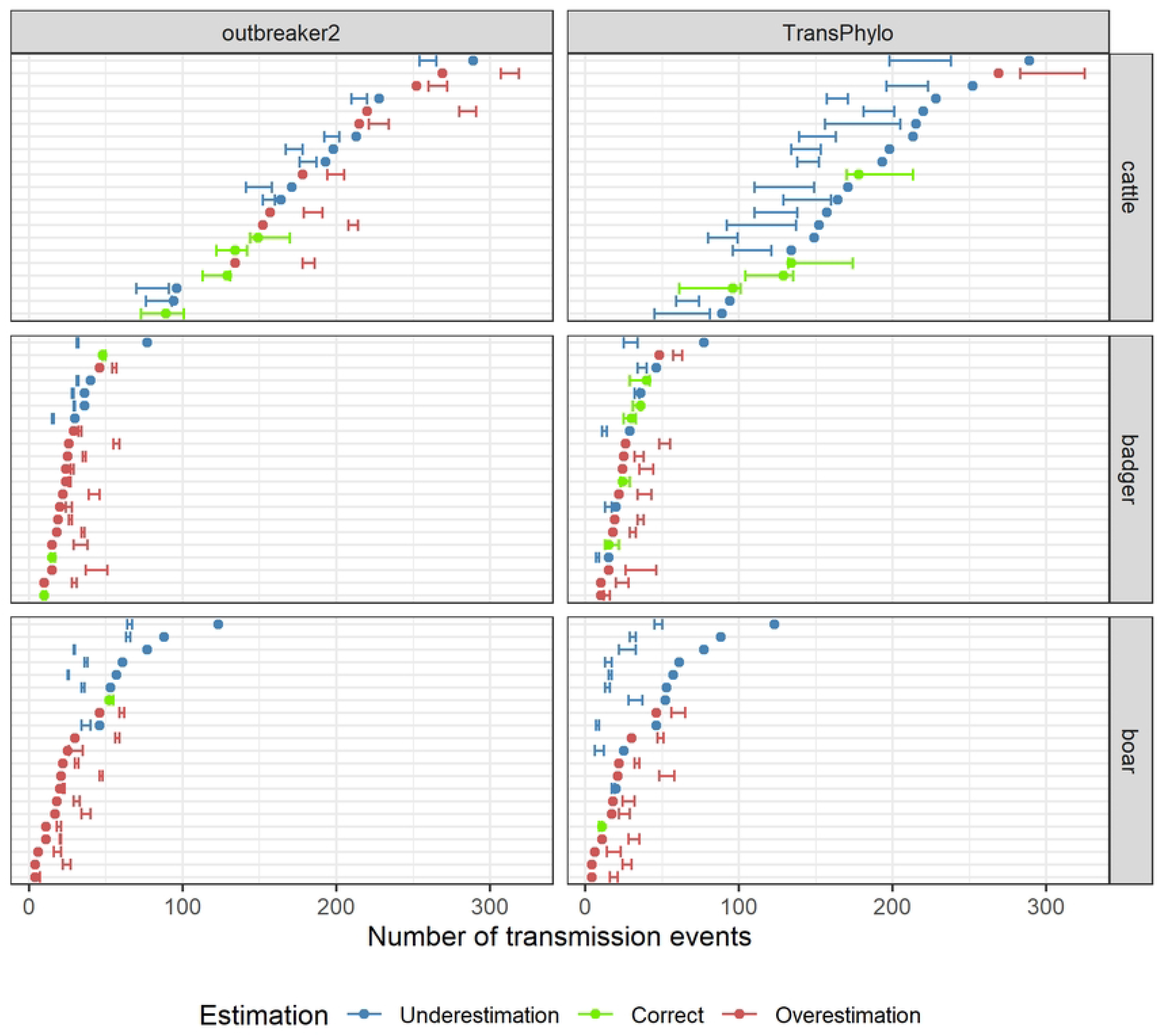
Credible interval of host-species contribution compared to simulated outbreaks in the high mutation rate scenario. The credible interval was either estimated by *outbreaker2* or by *TransPhylo*. The point corresponds to the number of transmission events due to each host-species in the simulated outbreak.

#### 3.2 Single-host system

Sequences simulated within a single-host system presented a lower proportion of unique sequences (median: 3.6%) and a lower mean transmission divergence (median: 0.14) (S7 Table).

Similarly to the multi-host systems, the accuracy was the highest for *TransPhylo* (6.5%), then *outbreaker2* (5.5%) and the lowest for seqTrack (2%) (S8-S9 Table). Super-spreaders were present in all trees reconstructed by seqTrack but also in trees reconstructed by *outbreaker2* (5/26 trees, median of maximum 39 transmission events due to a single super-spreader) and *TransPhylo* (10/26 trees, median: 39.5) (S10 Table).

The credible interval contained the simulated outbreak size in all 26 trees reconstructed by *outbreaker2* and in 16/26 trees reconstructed by *TransPhylo*, otherwise the outbreak size was overestimated (Fig 7 and S11 Table).

**Fig 7.**
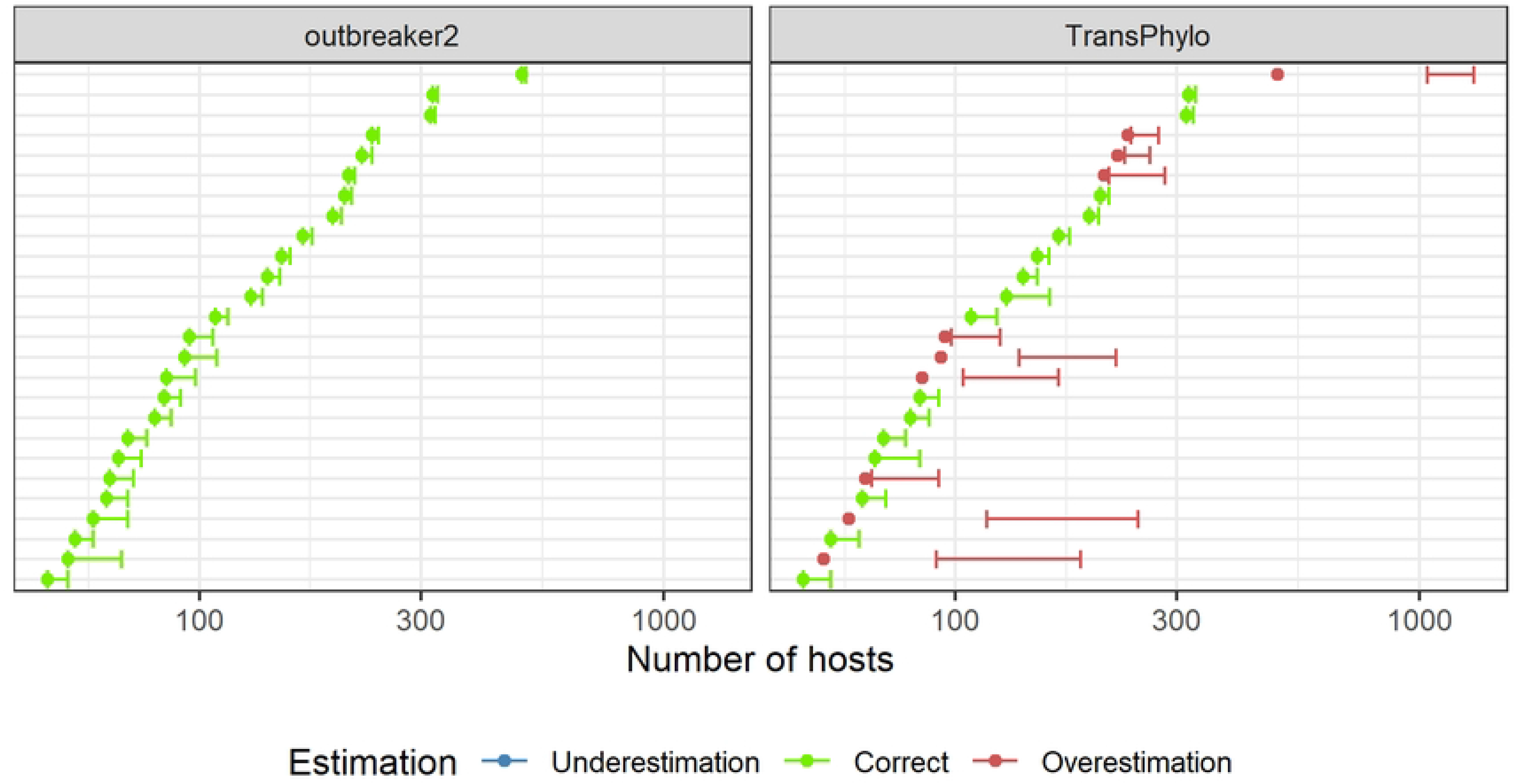
Outbreak size credible interval compared to simulated outbreak size in the single-host system scenario. The credible interval was either estimated by *outbreaker2* or by *TransPhylo*. The point corresponds to the simulated outbreak size.

#### 3.3 Dead-end epidemiological host

The credible interval never contained the simulated number of transmission events due to wild boars and wild boar contribution was overestimated in all 17 reconstructed trees (Fig 8). Otherwise, similarly to the multi-host systems without a dead-end epidemiological host, cattle contribution tended to be underestimated by both methods (17/17 trees for *outbreaker2* and 10/17 for *TransPhylo*) and badger contribution, overestimated by *outbreaker2* (15/17 trees in Fig 8, and S12 Table).

**Fig 8.**
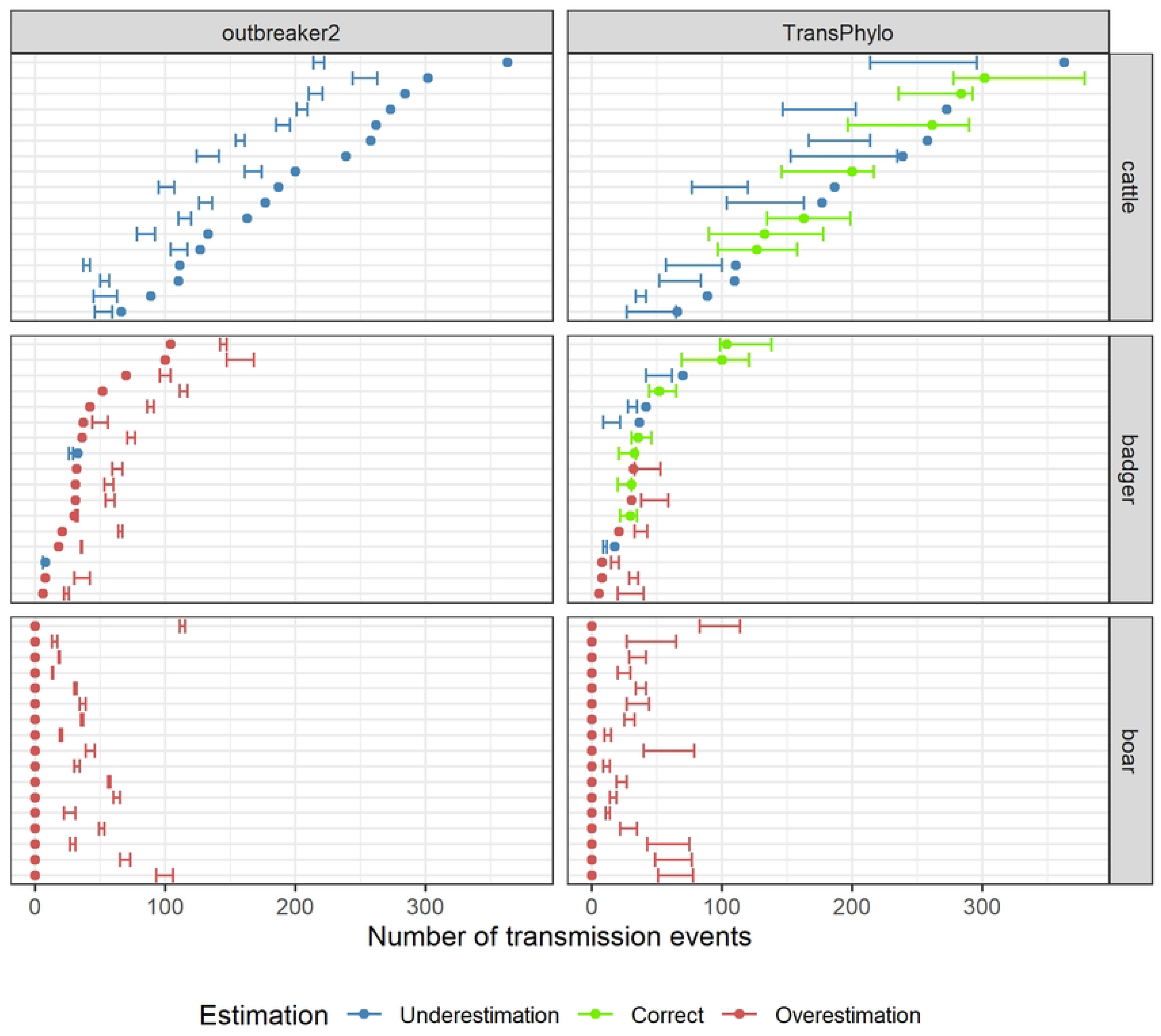
Credible interval of host-species contribution compared to simulated outbreaks in the dead-end epidemiological host scenario. The credible interval was either estimated by *outbreaker2* or by *TransPhylo*. The point corresponds to the number of transmission events due to each host-species in the simulated outbreak.

#### 3.4 Badger index

The proportion of correctly reconstructed badger index cases compared to cattle index cases was markedly lower for *outbreaker2* (28%), seqTrack (28%) and *TransPhylo* (11%).

For all transmission scenarios, even the reference multi-host scenario, similar results were obtained when considering the reference tree or the reconstructible outbreak (S2 Appendix and S11-S12 Tables).

## Discussion

In this work, we evaluated and compared the performances of three outbreak reconstruction methods on simulated *M. bovis* data in a multi-host system, as well as the impact of observation biases on these performances. *M. bovis*, characterized by a low mutation rate, is a prime example of a multi-host pathogen for which sampling biases complicate the estimation of host-species contribution to transmission, an estimation which is however necessary to select appropriate measures for disease control. Contrary to previous evaluations of outbreak reconstruction methods, the transmission model we used to simulate our data was not tailored to a specific method [25,31,52] but to the slowly evolving multi-host pathogen. Moreover, the epidemiological indicators we estimated were also relevant in a multi-host system and not just general performance indicators [53].

Reconstructing transmission trees can have multiple objectives according to the studied pathogen and epidemiological system, the most obvious objective is the accurate reconstruction of who-infected-whom. The proportion of correctly reconstructed transmission events (which we called accuracy) has previously been used to evaluate performances of outbreak reconstruction methods [25,53]. With the low mutation rate characteristic of *M. bovis*, we estimated poor accuracies (median accuracy lower than 9% for all three methods). Sobkowiak *et al.* compared these outbreak reconstruction methods on real *M. tuberculosis* data, which is also a slow-evolving pathogen, and estimated the positive predictive value (PPV), meaning the number of epidemiologically linked case-contact pairs that were correctly identified (preprint, [54]). Contrary to the accuracy indicator we estimated, the links between cases were not directed, we thus expected this study to estimate a higher number of correctly reconstructed cases. The PPV estimated by Sobkowiak *et al.* was 15% for *TransPhylo*, 11% for *outbreaker2* and 10% for seqTrack. These PPV values were in the range of values we estimated for accuracy and the ranking of methods was the same as the one we obtained (with *TransPhylo* as the best, followed by *outbreaker2*).

Accuracy was little influenced by the sampling biases or the complexity of the epidemiological system, however it was greatly dependent on the mutation rate. When the mutation rate was multiplied by a factor of 10 (∼6.6 x 10^-5^ substitutions per site per day), the accuracies we estimated more than doubled. In the study that presented and tested *outbreaker2*, Campbell *et al.* estimated the average proportion of transmission pairs correctly inferred when using solely temporal and genetic information from simulated Ebola virus (mutation rate: 0.31 x 10^-5^ per site per day) and SARS-CoV-1 (1.14 x 10^-5^ per site per day) outbreaks [25]. Moreover, Firestone *et al.* compared *TransPhylo* and *outbreaker2* on six FMDV outbreaks simulated with a high mutation rate (2.2 x 10^-5^ per site per day) and estimated the proportion of infected hosts (premises) for which the most likely source predicted was the true source [53]. Since both indicators corresponded to the accuracy we estimated, we expected similar results. However, Campbell *et al.* estimated an average accuracy of 29% (from the simulated Ebola data) and 70% (SARS-CoV-1). In addition, when genomic data was available for all infected hosts, the accuracy estimated by Firestone *et al.* was 4% for *TransPhylo* and 35% for *outbreaker2*. While these values were respectively higher for *outbreaker2* and lower for *TransPhylo* compared to the range of values we calculated, the ranking of methods obtained by Firestone *et al.* was the same as the one we obtained (with *outbreaker2* as the better of the two).

The lowest accuracy always being estimated for seqTrack could be due to the fact that this method does not consider a transmission model [21], but simply sampling dates and genetic distances. As mentioned by Nigsch *et al.*, seqTrack is thus strongly dependent on the temporal order of sampling dates and when the sampling order does not necessarily coincide with the infection order (here, because of imperfect case detection and sampling protocol varying according to host-species), “the order of ancestries cannot be inferred with certainty” [55]. Contrary to what we observed with *outbreaker2* and *TransPhylo*, trees reconstructed by seqTrack presented super-spreaders with extreme numbers of transmission events due to a single infected host (over a hundred transmissions) that lowered when considering a higher mutation rate. The low genetic diversity combined with the lack of a transmission model could therefore account for the reconstruction of super-spreaders, which in turn could contribute to the low accuracy. Similarly, the lower genetic diversity obtained with the single-host system could explain the presence of less prolific super-spreaders in trees reconstructed with *TransPhylo* and *outbreaker2*.

While we estimated poor accuracies for all three methods, a high proportion of correctly reconstructed directed transmission events is difficult to obtain and might not be the main objective when studying a multi-host system implicating wildlife or with a low sampling proportion. However, the presence of super-spreaders is an important indicator to consider since it highlighted the fact that seqTrack reconstructed unrealistic transmission dynamics with prolific super-spreaders.

Other than reconstructing who-infected-whom, outbreak reconstruction can aim to estimate epidemiological indicators, from which practical measures can be directly inferred. The first we studied was the outbreak size, which could by comparison with the number of sampled cases be informative *e.g.* of the need to increase the sampling effort [35]. Outbreak size estimation was sensitive to sampling biases, the complexity of the epidemiological system and also the mutation rate. The outbreak size was correctly estimated by *outbreaker2* but consistently overestimated by *TransPhylo*, even though we considered the same non-informative prior for the sampling proportion when implementing both methods. This overestimation could therefore be due to the fact that Didelot *et al.* developed this method to study partially sampled *M. tuberculosis* outbreaks and account for within-host diversity [31], whereas we assumed all cases sampled in the reference scheme and no within-host diversity in the sequence simulation. Furthermore, when not all sequences were sampled, better results were obtained for *TransPhylo* and the estimated outbreak size significantly lowered.

With a higher mutation rate, *TransPhylo* also overestimated the outbreak size, but to a lesser extent. Xu *et al.* developed in 2019 a method of simultaneous inference on multiple *M. tuberculosis* clusters based on *TransPhylo*. From this study, Xu *et al.* discussed the link between mutation rate and sampling proportion, explaining that with a faster assumed clock, the branches in the phylogenetic trees are shorter and *TransPhylo* is therefore less likely to place unsampled cases along them [56]. This could explain the lower effect we estimated.

Some epidemiological indicators are relevant only in the context of a multi-host system and reveal the host-species that should be primarily targeted, such as the identification of the host-species responsible for the outbreak and the accurate reconstruction of each host-species’ contribution to the outbreak. The index case indicator was sensitive to sampling biases and to the host-species of the index case. The proportion of correctly reconstructed host-species of the index case was high for *outbreaker2* and seqTrack (over 75%) when considering cattle index cases. However, the fact that *TransPhylo* could designate unsampled hosts as index cases, combined with a tendency to overestimate the outbreak size and thus, the number of unsampled hosts, could explain this method’s poorer performance. Moreover, biased sampling schemes generally led to a higher proportion of correctly reconstructed host-species of the index case, which could be explained by the fact that only non-index cases (wildlife) were concerned by these sampling schemes. Finally, this indicator was sensitive to the host-species responsible for the outbreak and had a poorer performance when the index case was a badger.

Host contribution estimation was influenced by the sampling biases and the complexity of the epidemiological system but not the mutation rate. With either mutation rates, *outbreaker2* and *TransPhylo* poorly reconstructed the contribution of each host-species and tended to underestimate the host-species that contributed the most to transmission (cattle) while overestimating those that contributed the least (wildlife). Both outbreak reconstruction methods were developed and tested on single-host systems [21,25,31], and not on multi-host systems where each host-species play a different role. While *TransPhylo* has previously been applied to multi-host systems, a human-deer SARS-CoV-2 system [35] and two badger-cattle bTB systems [57,58], the estimation of host-species contribution to transmission in these systems was not straightforward. The high number of unsampled cases estimated in the human-deer system (mean sampling proportion of 0.1%) complicated the inference of transmission events and while phylogenetic evidence seemed to support multiple human-to-deer spillover events, deer-to-human transmission could not be ruled out [35]. In a badger-cattle system in the South-West of England, van Tonder *et al.* were interested in between-species transmission and as such *TransPhylo* was implemented in addition to a Bayesian ancestral state reconstruction method (BASTA, [16]), which was primarily used to estimate the number of within-and between-species transitions [57]. Finally, Akhmetova *et al.* also implemented *TransPhylo* in addition to Bayesian phylogenetic methods in a badger-cattle system in Northern Ireland and highlighted a mostly cattle-driven (over 90% of strongly supported reconstructed transmission events) epidemic in the region [58].

Biases simulated with the sampling schemes resulted in a decrease in the number of infected hosts for which contribution estimates was overestimated. Therefore, when sampling schemes had a significant effect on host contribution, they tended to yield better results with this particular host-system and either lowered (for wildlife) or increased (for cattle) the estimated number of transmission events. In addition, neither *outbreaker2* nor *TransPhylo* could accurately reconstruct asymmetrical roles between host-species, *i.e.* the presence of a dead-end epidemiological host.

In the epidemiological multi-host system we extended, the basic reproduction number varied according to the combination of host-species considered [39]. Moreover, Bouchez-Zacria *et al.* calculated inter-and intra-species generation time distributions that showed a more rapid spread from cattle farms than from badger groups. We added to the transmission model, a third population of host-species (wild boars) that could transmit or not the pathogen. The complexity of this multi-host system could have contributed to the poor results we obtained for host-species contribution. Indeed, both *outbreaker2* and *TransPhylo* considered a single generation time, sampling time and/or offspring distribution for all three host-species, not accounting for host-species variation in the natural history of the disease nor the uneven transmission dynamics. A multi-host system where all three host-species contributed unevenly to transmission is not unusual, results obtained from Bayesian ancestral state reconstructions (Mascot, [19]) in other French regions point to the presence of similarly complex bTB multi-host systems [59]. Furthermore, the impact of said complexity on method performance does not only concern systems with multiple host-species, pathogens for which different categories (*e.g.* age groups and/or vaccination status [60]) of hosts can be defined (according to infectiousness or duration of infection) also constitute complex epidemiological systems. Finally, considering what can be reconstructed by the method (reconstructible outbreak) instead of the reference tree did not improve results for the outbreak size nor the host contribution indicators.

We were limited by practical considerations and the ensuing choices we made. With the *M. bovis* data we simulated, convergence was a limiting factor for *TransPhylo* but not for *outbreaker2*. Indeed, in order to limit the computational time, we fixed a maximum number of iterations, which narrowed the number of reconstructed trees we could compare to those that converged in less than 48 hours in BEAST2 and 12 hours in *TransPhylo*. Moreover, in order to better compare reconstructions, we used the same evolutionary model for the phylogenetic reconstruction in BEAST2. A more adapted evolutionary model could lead to a more accurate phylogenetic tree reconstruction and thus, a better performance from *TransPhylo*.

With the sequence simulation model we implemented, we simulated a low proportion of unique sequences from 13-year-long outbreaks, which is consistent with *M. bovis* low mutation rate. In the study on 167 *M. bovis* sequences in the South-West of France from which we selected the value of the mutation rate [12], the proportion of unique sequences isolated (37.1%) was around six times higher than the median proportion we simulated. The higher proportion of unique sequences in this previous study could be due to the fact that not all sequences are sampled in real data and that the outbreak lasted longer as suggested by the MRCA which was estimated to have been circulating 27 years earlier. In the sequence simulation model, we also considered the same mutation rate within all three host-species, however whether *M. bovis* evolves the same way within different host-species remains unknown. Indeed, *M. tuberculosis* mutation rates in humans may decrease during periods of latency, which differs from what was observed in non-human primates [61]. A similar phenomenon could lead to variability in the evolution of *M. bovis* within and between host-species [62] and thus, to the difference in the proportion of unique sequences observed and simulated.

We chose to compare results from the same epidemiological and genetic data for all three methods. While difficult to implement when studying a slowly evolving multi-host system that implicates wildlife, contact data can be directly incorporated in the transmission tree inference with *outbreaker2*. The addition of contact data led to higher accuracies than those obtained with only temporal and genetic data in simulated Ebola virus and SARS-CoV-1 outbreaks [25]. Similarly, when limited genetic diversity was expected in their study on *M. avium* ssp *paratuberculosis*, Nigsch *et al.* took advantage of the fact that seqTrack can incorporate additional data in the form of weighting matrices [55]. They thus resolved equally probable ancestries using known exposure time or susceptibility based on accepted epidemiological knowledge. Even when the method does not allow additional epidemiological data, Xu *et al.* mentioned that one of the strengths in their study on *M. tuberculosis* transmission within a Spanish cohort lied in the extensive contact investigation data that allowed them to validate the results of their genomic and *TransPhylo* analysis [63]. Using these available features and strategies could have improved results obtained for *outbreaker2* and seqTrack as well as help evaluate those from *TransPhylo*. However, since limited real contact data can be obtained for wildlife, we chose not to include additional epidemiological data in this study. Furthermore, we limited our study to only three methods, available in a package and that only needed sampling times as epidemiological data. Additional methods would be interesting to test on this simulated data, especially methods that simultaneously inferred phylogenetic and transmission trees like *phybreak* [26], since none were considered here. Finally, the simulated multi-host data could also be used to test Bayesian ancestral state reconstruction methods like Mascot [19], previously used to study complex bTB multi-host systems [59].

The overall poor performances we obtained for accuracy and host-species contribution, even without biased sampling schemes, suggest that when studying the transmission of a slowly evolving pathogen in complex multi-host systems, outbreak reconstruction methods should not be implemented alone but as a complement to epidemiological and phylogenetic methods. The difficulty in estimating host-species contribution highlights the need to develop new outbreak reconstruction methods adapted to complex epidemiological systems as well as evaluate these methods on data simulated in multi-host systems and not specific to the each method.

## Availability of data and materials

All simulated data and code used to simulate data, reconstruct outbreaks and evaluate methods are available on Github (https://github.com/duaulthel/bTBtreereconstruction.git).

## Acknowledgements

Not applicable.

## Supporting information

S1 Appendix. Details on transmission tree simulation, phylogenetic and transmission tree reconstruction.

S2 Appendix. Results on outbreak size and host contribution indicators using the reconstructible outbreak as a reference.

S1 Fig. Proportion of transmission pairs with 0, 1 and 2 SNPs between their sequences according to transmission scenario. Reference stands for the complex multi-host system where cattle are index cases and wild boars contribute to transmission. High mutation rate is the same scenario as the reference except for the higher mutation rate used to simulate sequences. Single-host stands for the only-cattle scenario. Dead-end host stands for the scenario where wild boars did not contribute to transmission and badger index, the reference scenario with badger as index cases.

S2 Fig. Credible interval of the number of transmission events due to cattle estimated by *outbreaker2* and *TransPhylo* compared (color) to the number in the simulated outbreak (point) according to sampling scheme. T stands for “temporal bias”, SB for “badger bias” and SW for “wild boar bias”. T+SB (T+SW) combined the temporal and the badger (wild boar) bias.

S3 Fig. Credible interval of the number of transmission events due to badgers estimated by *outbreaker2* and *TransPhylo* compared (color) to the number in the simulated outbreak (point) according to sampling scheme. T stands for “temporal bias”, SW for “wild boar bias” and T+SW combined the temporal and the wild boar bias.

S4 Fig. Credible interval of the number of transmission events due to wild boars estimated by *outbreaker2* and *TransPhylo* compared (color) to the number in the simulated outbreak (point) according to sampling scheme. T stands for “temporal bias”, SB for “badger bias” and T+SB combined the temporal and the badger bias.

S1 Table. Comparison between reference trees that converged in BEAST2 and *TransPhylo* and those that did not.

S2 Table. Proportion of reconstructed transmission events that were present in the reference trees according to method and sampling scheme.

S3 Table. Maximum number of transmission events a single super-spreader could be responsible for in a tree reconstructed by seqTrack and their host-species according to sampling scheme.

S4 Table. Proportion (%) of correctly reconstructed host-species of the index case according to method and sampling scheme.

S5 Table. Number of infected hosts present in the induced subtrees and reconstructed trees according to method and sampling scheme.

S6 Table. Number of transmission events due to each host-species in the reconstructible outbreak and reconstructed trees according to method and sampling scheme.

S7 Table. Comparison between reference trees that converged in BEAST2 and *TransPhylo* and those that did not, according to transmission scenario.

S8 Table. Proportion (%) of reconstructed transmission events that were present in the reference trees according to method and transmission scenario.

S9 Table. Accuracy tested with a Binomial GLM using method as the explanatory variable, according to transmission scenario.

S10 Table. Number of trees reconstructed where super-spreaders were present and the maximum number of transmission events a single super-spreader could be responsible for according to method and transmission scenario.

S11 Table. Outbreak size with a Negative Binomial GLM using method as the explanatory variable, according to transmission scenario.

S12 Table. Host contribution tested with a Negative Binomial GLM using method as the explanatory variable, according to transmission scenario.

